# The Sodium/Glucose Cotransporter 2 Inhibitor Empagliflozin Inhibits Long QT 3 Late Sodium Currents in a Mutation Specific Manner

**DOI:** 10.1101/2024.04.15.589662

**Authors:** Lynn C. Lunsonga, Mohammad Fatehi, Wentong Long, Amy J. Barr, Brittany Gruber, Arkapravo Chattopadhyay, Khaled Barakat, Andrew G. Edwards, Peter E. Light

## Abstract

**Background:** Sodium/glucose cotransporter 2 inhibitors (SGLT2is) such as empagliflozin have demonstrated substantial cardioprotective effects in patients with or without diabetes. The SGLT2is have been shown to selectively inhibit the late component of cardiac sodium current (late I_Na_). Induction of late I_Na_ is also the primary mechanism involved in the pathophysiology of congenital long QT syndrome type 3 (LQT3) gain-of-function mutations in the SCN5A gene that encodes the major cardiac sodium channel isoform Nav1.5. Therefore, we investigated the effect of empagliflozin on late I_Na_ in thirteen known LQT3 mutations located in distinct regions of the channel structure.

**Methods:** The whole-cell patch-clamp technique was used to investigate the effect of empagliflozin (10 µM) on late I_Na_ in recombinantly expressed Nav1.5 channels containing different LQT3 mutations. Molecular modeling of human Nav1.5 and simulations in a mathematical model of human ventricular myocytes were used to extrapolate our experimental results to excitation contraction coupling.

**Results:** Empagliflozin selectively inhibited late I_Na_ in LQT3 mutations residing in the inactivation gate region of Nav1.5, with no effect on either peak current or channel kinetics. In contrast, empagliflozin caused inhibition of both peak and late I_Na_ in mutations in the S4 voltage-sensing regions as well as changes in activation and inactivation kinetics and a slowing of recovery from inactivation. Empagliflozin had no effect on late/peak I_Na_ or channel kinetics in channels containing LQT3 mutations located in the putative empagliflozin binding region. Simulation of our experimental findings in a mathematical model of human ventricular myocytes predicts that empagliflozin may have a desirable therapeutic effect in LQT3 mutations located in the inactivation gate region.

**Conclusions:** Our results show that empagliflozin selectively inhibits late I_Na_, without affecting gating kinetics, in LQT3 mutations residing in the inactivation gate region. Patients with mutations in voltage-sensing regions are less suitable candidates as empagliflozin may prevent action potential firing. The SGLT2is may therefore be a promising novel precision medicine approach for patients with certain LQT3 mutations.

## Introduction

Voltage-gated sodium channels play an essential role in generating the cardiac action potential (AP) by producing a fast, but rapidly inactivating, inward sodium current (peak I_Na_) that generates the depolarizing upstroke of the AP. Under certain disease circumstances, including heart failure and ischemia reperfusion injury [1; 2], the likelihood of sodium channels failing to inactivate increases, leading to a persistent or late sodium current (late I_Na_) that results in AP prolongation. Ultimately, the increased duration of the AP causes early afterdepolarizations (EADs) that are a known electrophysiological substrate for the generation of life-threatening ventricular arrythmias [3; 4]. The induction of late I_Na_ also results in increased intracellular calcium levels through elevated reverse-mode Na^+^-Ca^2+^ exchange, adversely affecting calcium homeostasis and cardiac excitation contraction [5]. A sustained prolongation of the cardiac AP is reflected in the electrocardiogram (ECG) as an increase in QT interval, or long QT (LQT).

LQT syndrome can either be acquired, such as drug-induced [6], or congenital in nature [7]. LQT syndrome resulting from mutations in cardiac ion channels is the primary cause of sudden cardiac death (SCD) in patients below the age of 35 [8]. Gain-of-function LQT3 mutations are found in the *SCN5A* gene that encodes the major cardiac sodium channel isoform Nav1.5. Although LQT3 mutations account for only 5-10% of total LQT syndrome cases, it has the most malignant LQT phenotype with the fewest treatment options, and a 10-year survival rate of <50% [9; 10]. Whereas LQT1 and LQT2 syndrome are caused by loss-of-function mutations in potassium channels and can be managed with β-blockers and lifestyle modifications, treatment options and prognosis for LQT3 are still unfavorable [11]. Most patients are asymptomatic and are diagnosed by genetic screening for known mutations, usually in response to a family history of LQT, or coincidental findings of prolonged QT interval on their ECG. Clinical presentation can include syncope, seizures, and SCD in severe cases. Most symptoms occur during rest and are bradycardia-triggered, as QT-prolongation is most evident at lower heart rates in LQT3 [8; 12]. As a result, β-blockers are generally ineffective and prophylactic placement of implantable cardioverter defibrillators (ICDs) is required in the majority of LQT3 patients [13], albeit with significant drawbacks associated with ICD usage. Therefore, there is an urgent need to identify novel therapeutics that specifically target late I_Na_ in patients harbouring LQT3 mutations.

In this regard, the sodium/glucose cotransporter 2 inhibitors (SGLT2is) may represent an ideal candidate for further exploration as a therapy for LQT3. The first generation of SGLT2is were FDA-approved in 2013/2014 for the treatment of hyperglycemia in type 2 diabetes patients. Surprisingly, large-scale cardiovascular safety trials demonstrated marked cardiovascular benefits of SGLT2is compared to placebo. EMPA-REG OUTCOME (Empagliflozin, Cardiovascular Outcomes, and Mortality in Type 2 Diabetes), CANVAS (Canagliflozin Cardiovascular Assessment Study), and DECLARE-TIMI 58 (Dapagliflozin and Cardiovascular Outcomes in Type 2 Diabetes), have shown that patients with type 2 diabetes who received SGLT2is had a 30-40% relative risk reduction in death from cardiovascular causes, all-cause mortality, and hospitalization for heart failure [14–17]. The EMPEROR-Reduced (Empagliflozin Outcome Trial in Patients with Chronic Heart Failure and a Reduced Ejection Fraction), EMPEROR-Preserved (Empagliflozin in Heart Failure with a Preserved Ejection Fraction), and DAPA-HF (Dapagliflozin in Patients with Heart Failure and Reduced Ejection Fraction) trials subsequently demonstrated a cardiovascular benefit of SGLT2is in the absence of diabetes [18–20]. Interestingly, the DAPA-HF trial data suggest that dapagliflozin also seems to have an anti-arrhythmic effect [21; 22].

To date, the exact mechanism by which SGLT2is exert their cardioprotective properties is a topic of intensive investigation. Canonically, SGLT2is exert their hypoglycemic effect by inhibiting SGLT2 transporters in the proximal tubule of the kidneys, resulting in a reduction of renal glucose reabsorption [23]. It is well-established that SGLT2is have hemodynamic effects like decreasing blood pressure and plasma volume through natriuresis. However, recent studies have suggested a high probability of a direct effect of SGLT2is on cardiac muscle [24]. Various cardiac targets have been proposed to be affected by SGLT2is, including the NLRP3 inflammasome, calcium/calmodulin-dependent protein kinase II (CaMKII), and selective inhibition of late I_Na_ by our group [25–27].

This unexpected inhibitory effect of SGLT2is on cardiac late I_Na_ suggests that this class of drug may be a novel and timely approach to effectively treat patients with LQT3 mutations. In this regard, our group recently demonstrated late I_Na_ inhibitory effects of empagliflozin in two common LQT3 mutations [27]. Therefore, the objective of this current study was to comprehensively characterize the effects of empagliflozin on Nav1.5 currents containing 13 different LQT3 mutations located in distinct regions of the channel structure.

## Methods

### Cell Culture

HEK293T cells were grown in T25 cell culture flasks with a medium composed of Dulbecco’s Modified Eagle’s Medium (Sigma, USA), 7.5% fetal bovine serum, and 1% penicillin-streptomycin 10,000 U/mL (Cytiva, USA). The cells were kept in an incubator at 37 °C with 5% CO_2_ and split when 80-90% confluency was achieved. Splitting was accomplished by trypsinization with Trypsin-EDTA 0.25%, followed by resuspension and neutralization in cell medium. Cells were seeded on coverslips in 35 mm petri dishes at ∼70% confluency for experimental use.

### Nav1.5 Site-Directed Mutagenesis and Transfection in HEK293T Cells

LQT3 mutations were engineered into human wild-type (WT) Nav1.5 gene (*SCN5A*) in pcDNA3.1(+)/C-(K)DYK vector by overlap extension, using the Q5^®^ High-fidelity 2x Master Mix (New England Biolabs, USA). Melting temperatures (T_m_) were calculated by using the New England Biolabs calculator [22]. PCR cycles were initiated at 98 °C for 30 seconds to denature the template DNA, followed by 18 amplification cycles. Each amplification cycle consisted of 98 °C for 10 seconds, T_m_ of primer-template part minus 5 °C for 30 seconds, and 72 °C for 10 minutes (about 1 minute per 1 kbp of the template DNA). Amplification cycles were followed by an annealing step at T_m_ of primer-primer part minus 5 °C for 1 minute, and a final extension at 72 °C for 30 minutes. Amino acid deletions for ΔKPQ were generated by use of the QuikChange Lightning Site-Directed Mutagenesis kit (Agilent, USA). PCR products were cloned into XL-10 Gold Ultracompetent cells. DNA was isolated and purified by GeneElute^TM^ Plasmid Miniprep kits (Sigma, USA). Presence of the mutations was verified by DNA sequencing (The Applied Genomics Core, University of Alberta, Canada). The primer sequences used to generate the LQT3 mutants in Nav1.5 are listed in table 1. R1623Q primer sequences were obtained from a previous study [28]. LQT3 mutations were co-transfected with a green fluorescent protein (GFP) vector into HEK293T cells, using X-tremeGENE Reagent (Sigma, USA) with a 1:1.5 ratio for DNA and X-tremeGENE reagent. Transfected cells were seated on coverslips and were available for whole-cell patch-clamp 48-72 hours after transfection.

**Table 1.**
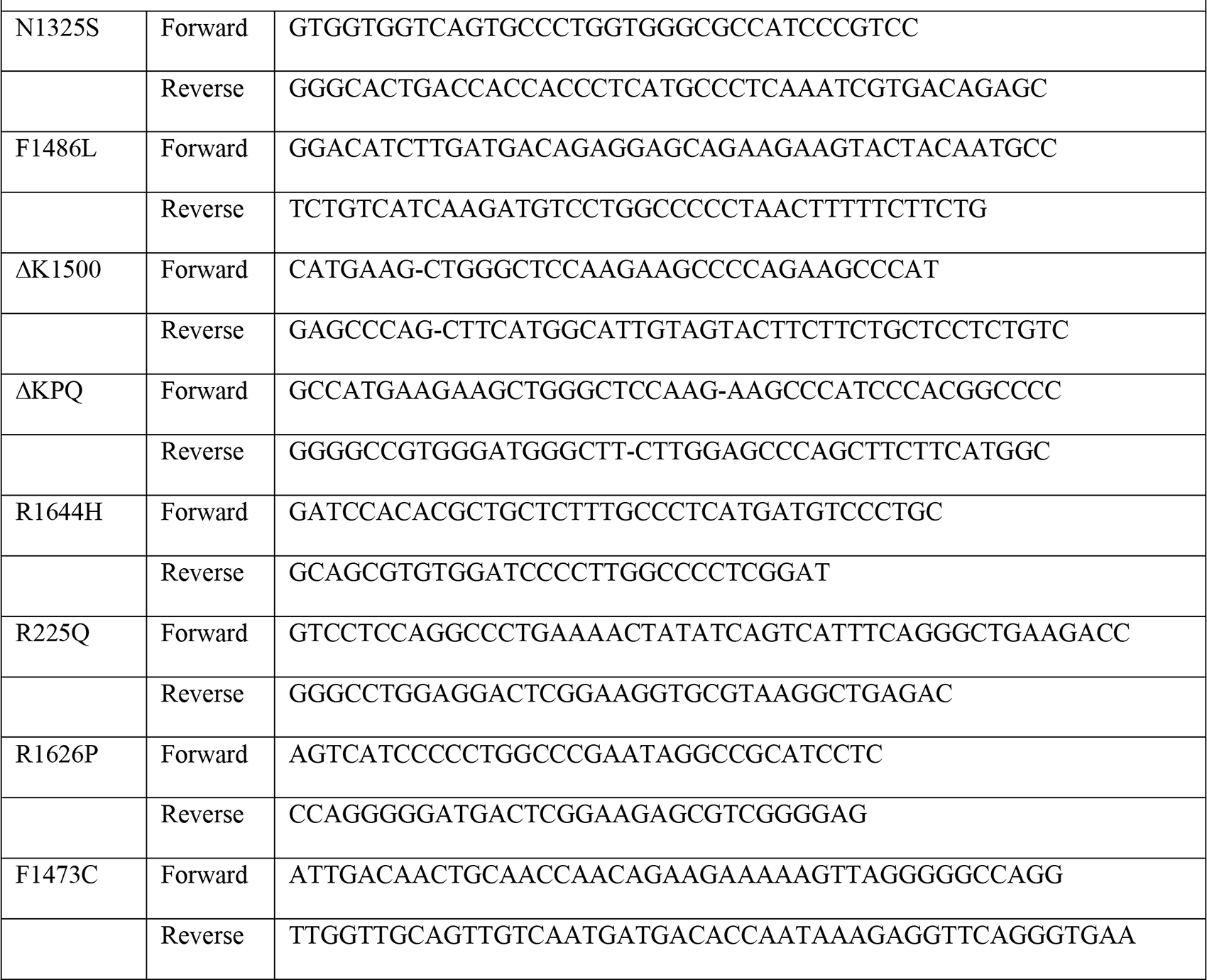
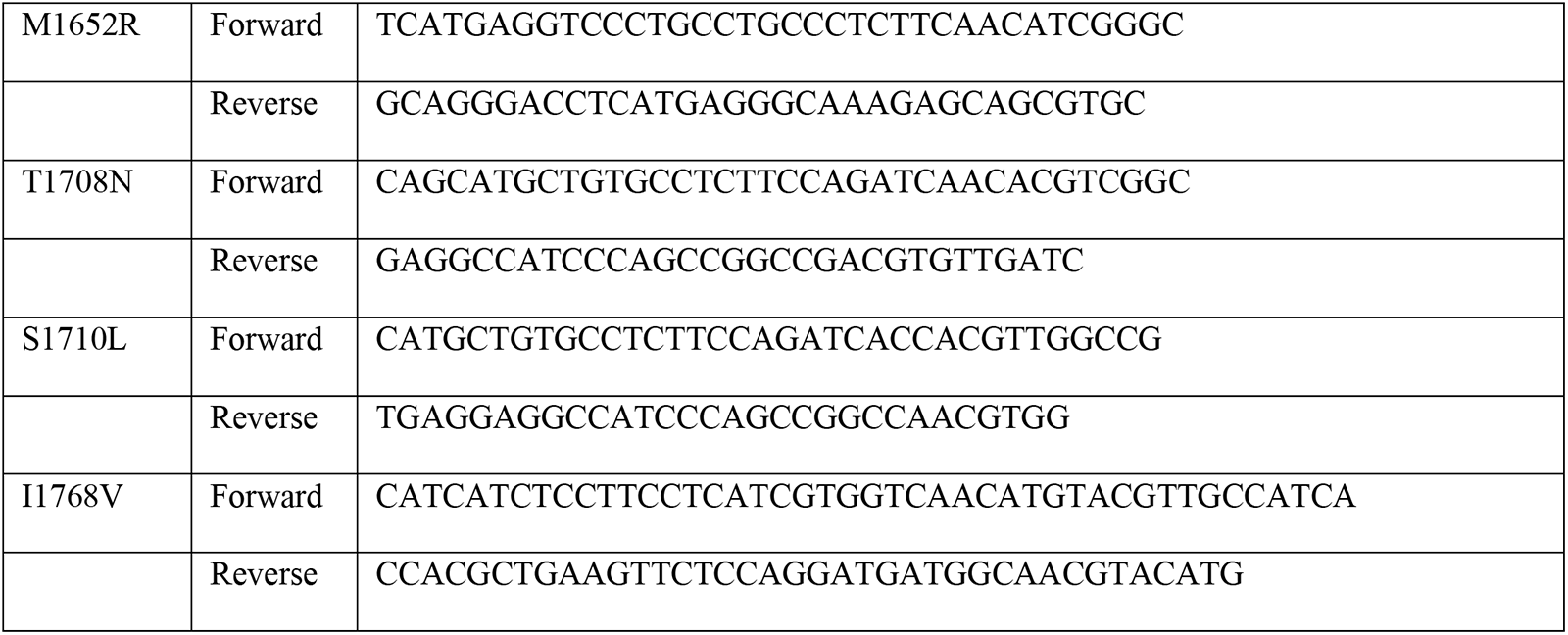
Nav1.5 Mutants Primer Sequences.

### Nav1.5 Current Recording

The whole-cell patch-clamp technique was used to investigate sodium currents in HEK293T cells, co-transfected with expression vectors for GFP and mutant human Nav1.5. Sodium currents from GFP-positive cells were recorded at room temperature (20-22 °C) 48-72 hours after transfection. Pipettes were pulled from borosilicate thin wall glass (Warner Instruments, USA) using a P-87 micropipette puller (Sutter Instruments, Novato, CA), after which the tips were fire-polished on a microforge to yield pipette resistances of 3-5 MΩ after filling with intracellular pipette solution. Pipette solution contained (in mmol/L) 138 CsCl, 12 HEPES, 11 glucose (C_6_H_12_O_6_), 1 MgCl_2_, and 1 CaCl_2_ (pH was adjusted to 7.2 with CsOH). Cells were bathed in extracellular solution, containing (in mmol/L) 140 NaCl, 2 MgCl_2_-6H_2_O, 2 CaCl_2_, 3 KCl, 10 HEPES, 5 tetraethylammonium bromide (to inhibit endogenous voltage-gated potassium channels in HEK293T cells). pH was adjusted to 7.4 with NaOH. After establishing a giga seal, the whole-cell configuration of patch-clamp technique was achieved by gentle suction. Whole-cell sodium currents were recorded by stepping from a holding potential of -130 to 20 mV for 400 ms by 10 mV incremental steps. Time-dependent recovery from inactivation was measured by peak currents P2 (20 ms at 0 mV) at various time intervals (Δt 1-100 ms) after P1, holding at -80 mV. Empagliflozin 10 µM was added to the extracellular solution after obtaining the initial control recording with dimethyl sulfoxide (DMSO, 0.1%) as vehicle, via a multi-channel perfusion system. Peak and late I_Na_ were recorded, digitized, and stored using an Axopatch 200B amplifier, Digidata 1322A, and pClamp10.2 (Molecular Devices, Union City, CA) [27].

### Homology Modeling of Human Nav1.5 Channel

The two cryoEM structures of the Nav1.5 channel (PDB entries 7DTC and 6LQA) were used as a starting point to build the full-length structure of the channel. While these two available structures encompass most of the structural regions of the channel (i.e., complete transmembrane domain, the extracellular segments, and the III-IV linker), they both lack several intracellular segments, including CaMKII phosphorylation sites. To construct and model these missing loops, we utilized the SWISS-MODEL server by providing the 7DTC and 6LQA structures as templates. The sequence of the full-length Nav1.5 protein was downloaded from the UniprotKB database (access code: Q14524, SCN5A_HUMAN) and was supplied to the SWISS-MODEL server. It is important to emphasize that our final model represents the human Nav1.5 channel in its closed state and consists of 4 transmembrane domains: DI (127-416), DII (712-939), DIII (1201-1470), and DIV (1524-1772). The cytoplasmic regions of Nav1.5 were not included due to lacking a good template. The generated model was further used as a starting point to generate *in silico* mutational analysis as indicated in figures 2-4. In generating these mutations, we utilized the molecular operating environment (MOE) software, by changing each residue to its corresponding mutation, while exploring the rotamers of the side chains to avoid any potential steric clashes with the surrounding residues.

### Mathematical Modeling of Nav1.5 Gating and Human Ventricular Myocyte Action Potential

To simulate the functional implications of empagliflozin on non-diseased and LQT3 ventricular APs, we first generated mathematical models of WT, ΔKPQ (representative class I) and R225Q (representative class II) Nav1.5 in the presence and absence of empagliflozin (five I_Na_ models in total). These were developed by refitting the cardiac I_Na_ model published by Clancy *et al*. [29] to the data presented herein and in Philippaert *et al* [27]. The resulting models were embedded in the Tomek 2019 human ventricular myocyte model (ToR-ORd, [30]) and paced to steady state (500 beats) at 1 Hz. The final 10s was used for visualization. Further details of the mathematical modeling and code can be found in the supplemental methods S1.

### Data and Statistical Analysis

Statistical analysis was performed using Clampfit 10.2 and GraphPad Prism 10.0.2. Curve fitting was obtained using Boltzmann equation. All values are presented as mean ± SEM. Unpaired T-tests and one-way ANOVA followed by Dunnett’s post hoc test were performed to determine significance. Statistical significances are indicated as not significant (ns, P>0.05), * (P<0.05), ** (P<0.01), and *** (P<0.001). Structural model figures were generated using the Visual Molecular Dynamics (VMD) software.

## Results

### LQT3 Mutations

The pore-forming α-subunit of Nav1.5 is encoded by the *SCN5A* gene and located on chromosome 3p21. It consists of four homologous transmembrane domains (DI-DIV), linked to each other through intracellular loops (figure 1). Every domain consists of six transmembrane segments (S1-S6), of which S4 is positively charged and involved in voltage-dependent channel activation. Inactivation of the channel, generated by the intracellular loop between DIII and DIV, leaves the channel refractory after repolarization of the AP, before the channel recovers to its closed state. The actual sodium conducting channel pores are located between S5 and S6 of each domain [3; 31; 32]. The vast majority of LQT3 cases are caused by autosomal dominant mutations in *SCN5A*, and they can be categorized into different classes based on their location within the channel. We designated class I mutations as those involved in inactivation gating between DIII and IV, and include N1325S, F1486L, ΔK1500, ΔKPQ (in-frame deletion of K1505, P1506, and Q1507), and R1644H [33–45]. Class II mutations were designated as those residing in the voltage-sensing regions (S4 activation helices), which include R225Q, R1623Q, and R1626P [9; 46]. We designated class III mutations as those that are in the putative SGLT2i binding site (F1473C, M1652R, T1708N, S1710L, and I1768V) [47–52], previously described by our group [27].

**Figure 1.**
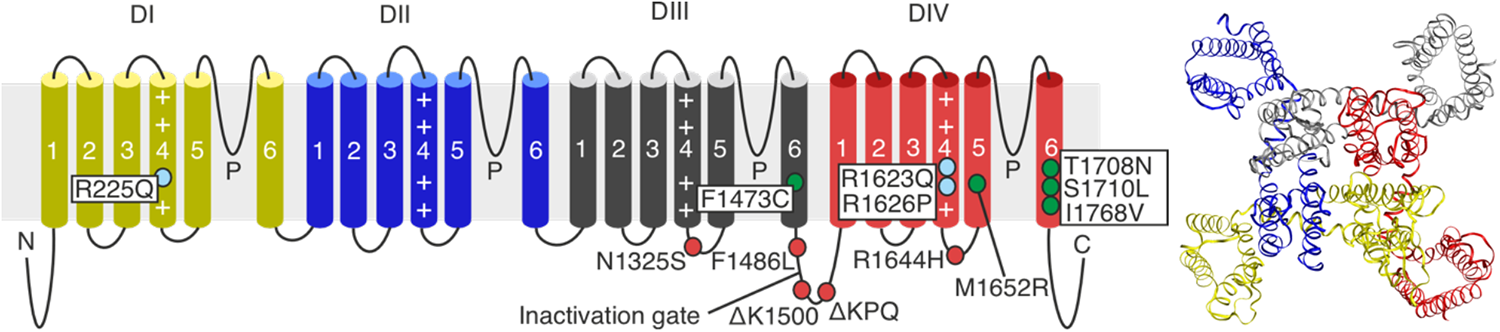
Schematic illustration of the structure of cardiac sodium channel Nav1.5 α-subunit. Locations of class I (red), class II (light blue), and class III (green) long QT syndrome type 3 mutations (left). Structural model top view of wildtype Nav1.5 (right). *P = pore loops*

### Empagliflozin Inhibits Late I_Na_ in Sodium Channels Carrying Long QT Syndrome 3 Mutations Residing in the Inactivation Gate Region

The inactivation gate region of the sodium channel is responsible for the rapid inactivation of the channel. It is therefore a common region in which to find LQT3 mutations that interfere with the appropriate fast inactivation process, leading to the development of late I_Na_. Typically, mutations in this region permit a fast initial inactivation but prevent the inactivation gate from closing completely, such that a small persistent plateau current is observed over several hundred milliseconds (Figure 1.). As induction of late I_Na_ is the primary pathophysiological mechanism of LQT3 syndrome, we explored the effect of empagliflozin on cardiac sodium currents of Nav1.5 channels containing LQT3 mutations in the inactivation gate region. [53]. Mutant human Nav1.5 currents containing either the N1325S, F1486L, ΔK1500, ΔKPQ (in-frame deletion of K1505/P1506/Q1507) or R1644H [33–45] where measure before and after the application of empagliflozin (10 μM). Empagliflozin inhibited late I_Na_ in all these class I LQT3 mutants residing in the inactivation gate (Figure 2). Peak I_Na_, inactivation, and activation were unaltered, indicating that empagliflozin selectively inhibits late I_Na_ in these mutants without affecting channel activation/inactivation kinetics.

**Figure 2.**
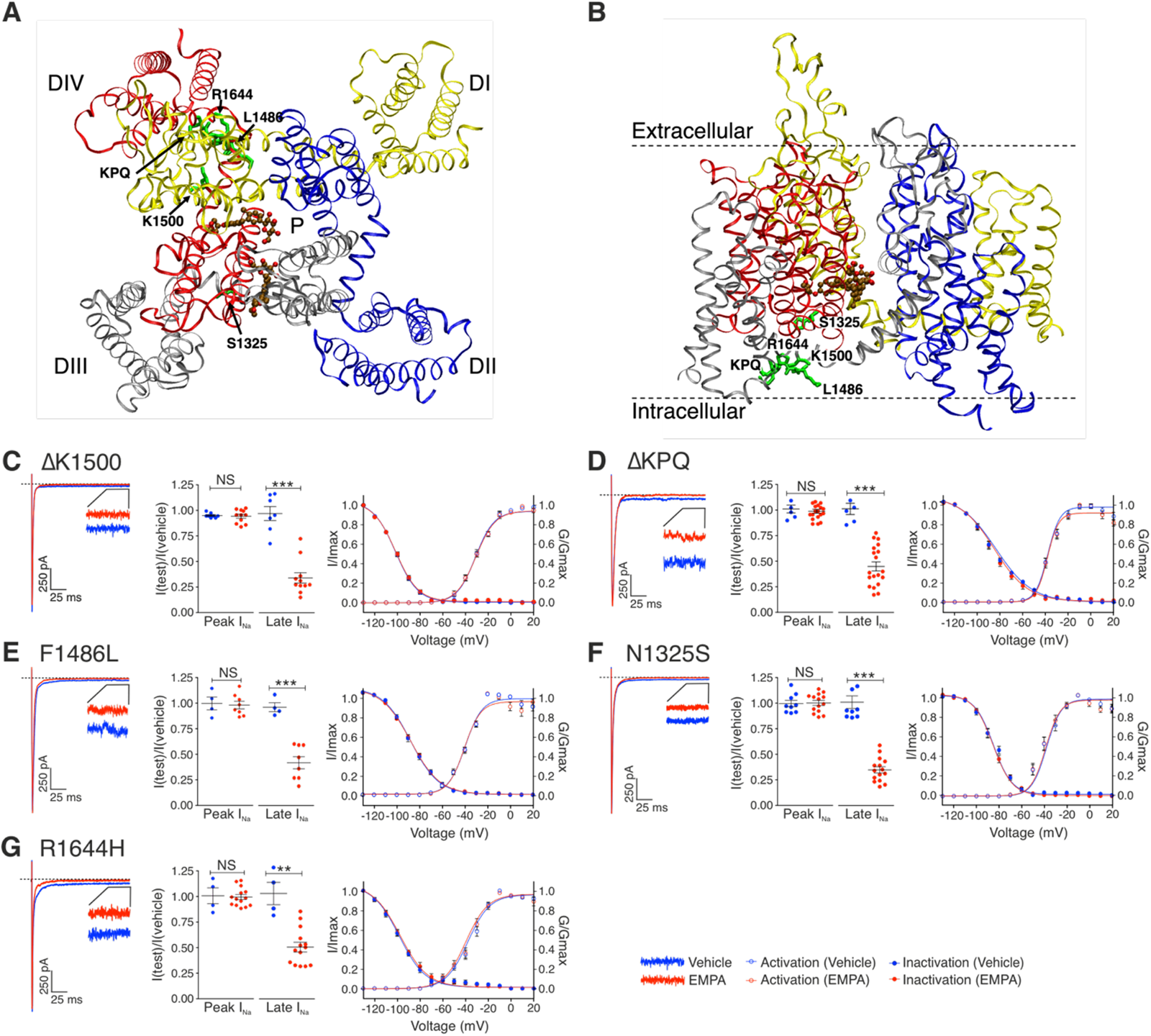
Empagliflozin inhibits the late component of cardiac sodium channel current in long QT syndrome type 3 mutations residing in the inactivation gate of Nav1.5. Structural modeling of human Nav1.5 with class I long QT syndrome type 3 amino acid residues in the inactivation gate region from an extracellular (A) and transmembrane (B) viewpoint. Docking of empagliflozin structure (brown) to the fDI-DIV and fDIII-IV sites is depicted, and residues are visible as green sticks. (C-G) Representative whole-cell patch-clamp recordings (left panels) showing the effects of empagliflozin on sodium currents from single class I mutated HEK293T cells (centre panels). Grouped data showing the effects of empagliflozin on channel activation and inactivation (right panels). *Grouped data are from 4-19 cell recordings per group. Data are presented as means ± SEM; *p<0.05; **p<0.01; ***p<0.001. I/Imax indicates test current/maximum current; G/Gmax indicates test conductance/maximum conductance*.

### Empagliflozin Inhibits Both Peak and Late I_Na_ in Long QT Syndrome 3 Mutations Residing in the S4 Voltage-Sensing Regions

Upon repolarization of the membrane potential, positively charged amino acid residues (mostly arginine and lysine) in the S4 segment of each domain initiate a conformational change, affecting the activation inactivation processes. S4 segments in domains I, II, and III are generally more involved in channel activation, whereas S4 in domain IV is associated with channel inactivation [54]. The functional variance arises from differences in specific amino acid residues and interactions with surrounding residues in neighboring segments. LQT3 mutations residing in the S4 regions that replace a positively charged residues with an uncharged one, induce late I_Na_ by altering these kinetic processes. We designated three of these S4 LQT3 mutations (R225Q, R1623Q, and R1626P [9; 46]) as Class II mutations. These mutations, in contrast to Class I mutations in the inactivation gate region, displayed a pronounced slowing of the of the initial fast inactivation process (Figure 3) as well as the development of late I_Na_. The addition of empagliflozin inhibited late I_Na_ in all Class II mutations but also significantly inhibited peak I_Na_ (Figure 3). With respect to activation and inactivation kinetics, R225Q channel inactivation was shifted to more negative potentials by empagliflozin, whilst activation was shifted to more positive potentials (Figure 3C). Consequently, this reduction of the window current, indicating the overlap of voltage-dependence of activation and inactivation, reduces the probability of spontaneous reopening of non-inactivated channels. In R1626P and R1623Q channels, empagliflozin similarly caused a negative shift of inactivation, whereas activation remained unaltered (figure 3D and 3E). These results are in accordance with previous research showing a preferential role of S4 in domain DIV in fast inactivation [54]. Empagliflozin-induced reduction in peak I_Na_ is likely resulting from the negative shift in inactivation, as more negative voltages are indicating are required to allow channels to fully recover from inactivation. In other words, by shifting the half-inactivation voltage to more negative potentials, empagliflozin increases the proportion of channels in the inactivated state and thus reduces peak I_Na_ through a decrease in fraction of channels available to open. As the upstroke of the cardiac AP is reliant on a fast inward sodium current, it is expected that any reduction in peak I_Na_ may prevent AP firing. These results suggest that empagliflozin would be an unfavorable choice of therapeutic for patients carrying these class II LQT3 mutations due to excessive inhibition of peak I_Na_.

**Figure 3.**
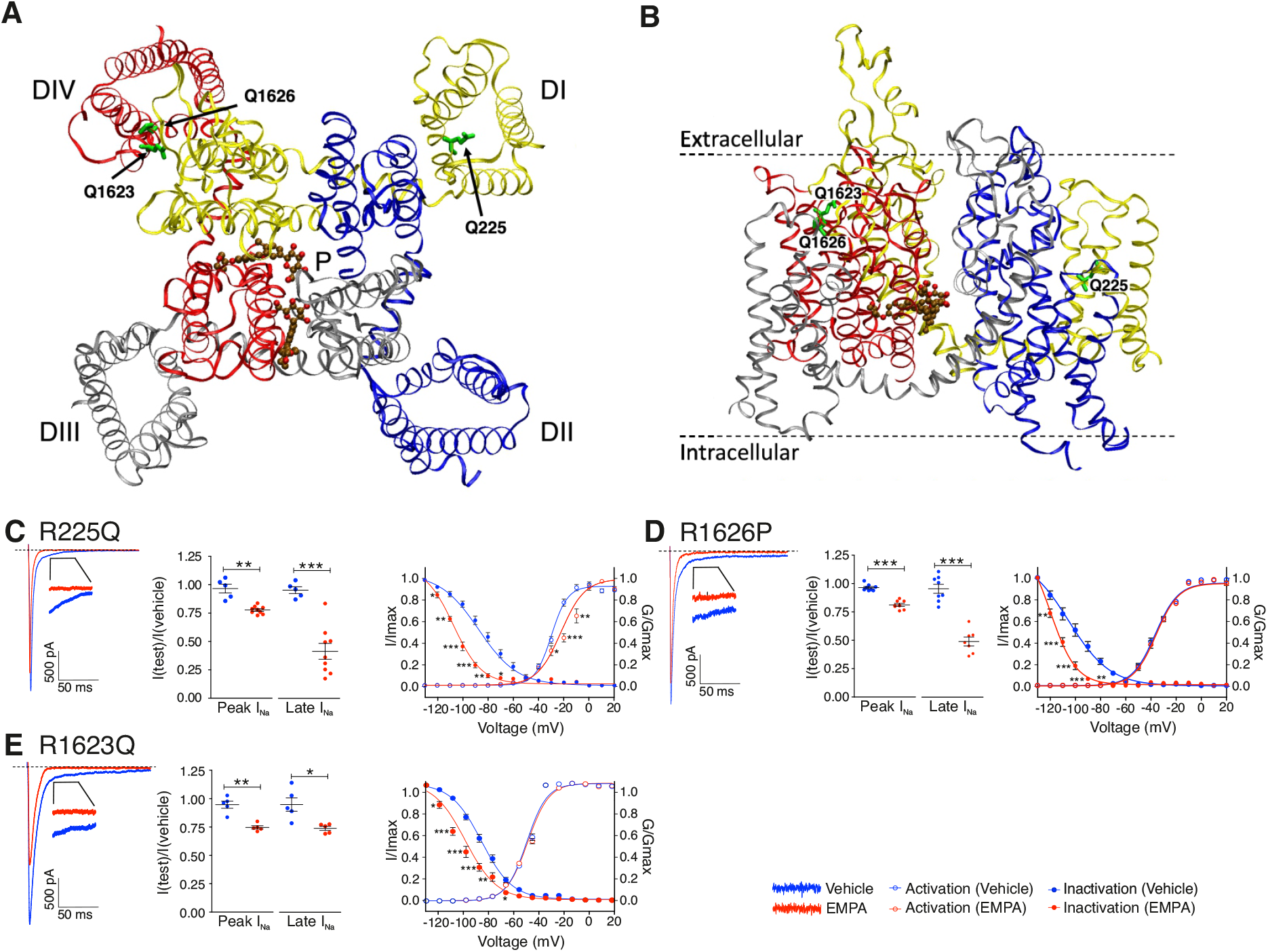
Empagliflozin inhibits both peak and late sodium currents of Nav1.5 channels with long QT syndrome type 3 mutations residing in the S4 voltage-sensing regions. Structural modeling of human Nav1.5 with class II long QT syndrome type 3 amino acid residues in the S4 voltage-sensing region from an extracellular (A) and transmembrane (B) viewpoint. Docking of empagliflozin structure (brown) to the fDI-DIV and fDIII-IV sites is depicted, and residues are visible as green sticks. (C-G) Representative whole-cell patch-clamp recordings (left panels) showing the effects of empagliflozin on sodium currents from single class II mutated HEK293T cells (centre panels). Grouped data showing the effects of empagliflozin on channel activation and inactivation (right panels). *Grouped data are from 4-9 cell recordings per group. Data are presented as means ± SEM; *p<0.05; **p<0.01; ***p<0.001. I/Imax indicates test current/maximum current; G/Gmax indicates test conductance/maximum conductance*.

### Empagliflozin Does Not Inhibit Late I_Na_ in Long QT Syndrome 3 Mutations Residing in the Putative Empagliflozin Binding Pocket

Our previous research investigated possible binding sites of empagliflozin within Nav1.5 [27]. *In silico* docking of empagliflozin to a three-dimensional homology model of open state human Nav1.5 allowed for molecular dynamic simulation. Two putative binding sites for empagliflozin were predicted at fenestration sites fDI-DIV and fDIII-IV, between DI-DIV and DIII-IV, respectively. The latter is a commonly known binding region for sodium channel inhibitors such as lidocaine [55] and our own point-mutational studies also support DIII-IV as a major binding region for empagliflozin. Therefore, we identified five LQT3 mutations that reside in this region (F1473C, M1652R, T1708N, S1710L, and I1768V) and tested the effects of empagliflozin in Nav1.5 channels containing each of these mutations [47–52] (Figure 4A and 4B).

Interestingly, empagliflozin did not inhibit late I_Na_ in any of these mutations residing in the predicted binding pocket, indicating possible interference of these mutations with empagliflozin binding and consequent lacking inhibitory effect on late I_Na_ (figure 4C-G). It is tempting to speculate that all tested Class III mutation sites in this study are involved in establishment of a stable interaction between Nav1.5 and empagliflozin. It is therefore predicted that patients with mutations in the putative SGLT2i binding site are likely to be SGLT2i-insensitive.

### Empagliflozin Inhibits Slowing of Recovery from Inactivation Only in Long QT Syndrome 3 Mutants Residing in S4 Voltage-Sensing Regions

As gain-of-function mutations of the Nav1.5 channel cause an increase late I_Na_, one mechanism of reducing late I_Na_ is by stabilizing the inactivated state of the channel and this was observed in the Class II mutants. As a consequence of inactivation state stabilization, it is predicted that a time-dependent slowing of recovery from inactivation may be observed in Class II LQT3 mutants. Accordingly, a protocol for time-dependent recovery was conducted in WT, ΔK1500 (Class I) and R225Q (Class II). Cells were voltage-clamped at -80 mV, and a first pulse (P1) at 0 mV was followed by a second pulse (P2) at increasingly longer time intervals (Δt from 1-100 ms). As predicted, only the R225Q Class II mutant channel, that previously showed a negative shift of inactivation in the presence of empagliflozin, displayed slowing of recovery from inactivation by empagliflozin (Figure 5).

**Figure 4.**
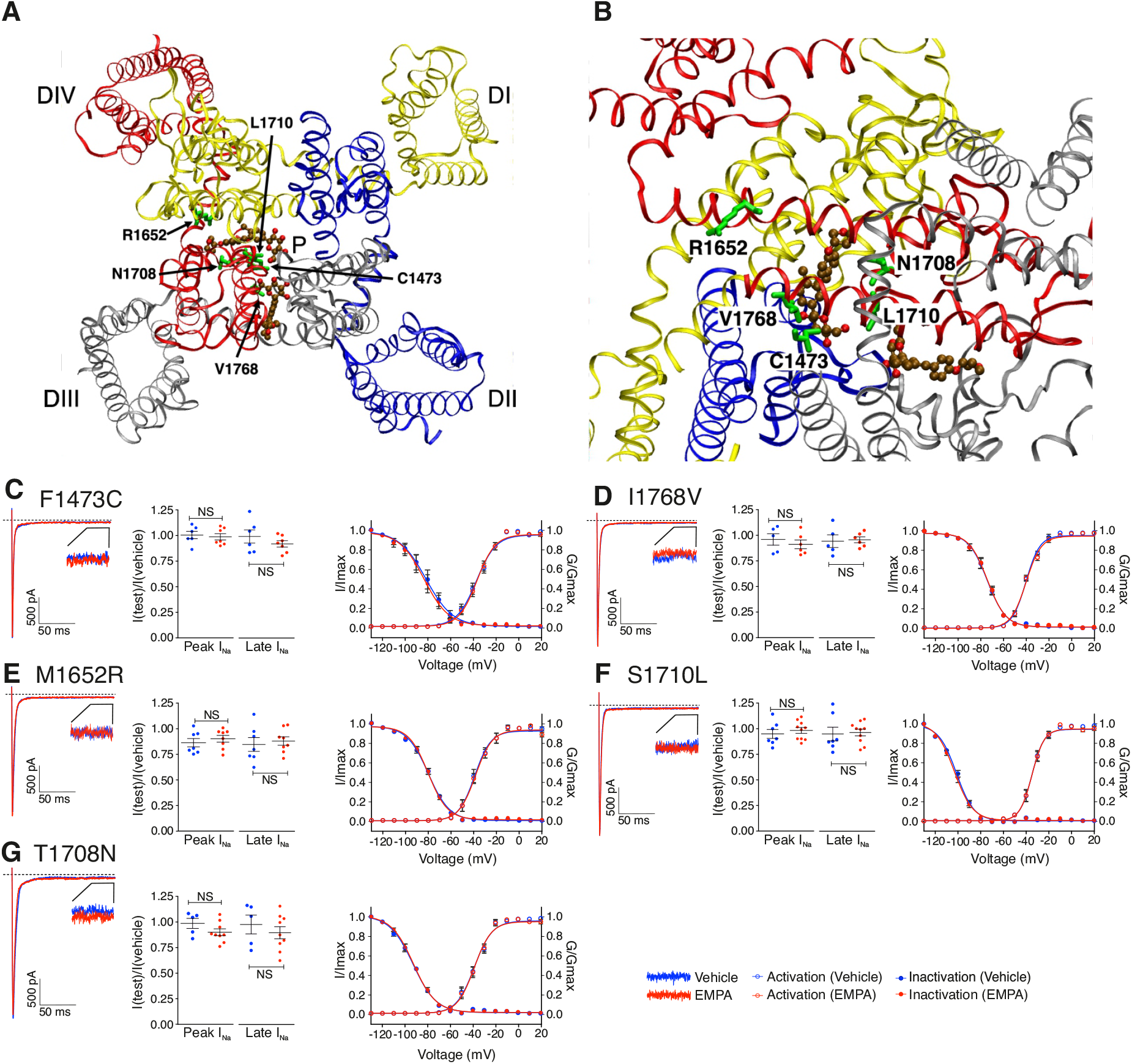
Empagliflozin does not inhibit late sodium currents of Nav1.5 channels with long QT syndrome type 3 mutations residing in the putative SGLT2i binding pocket. Structural modeling of human Nav1.5 with class III long QT syndrome type 3 amino acid residues in the putative empagliflozin binding region from an extracellular (A) and enlarged (B) viewpoint. Docking of empagliflozin structure (brown) to the fDI-DIV and fDIII-IV sites is depicted, and residues are visible as green sticks. (C-G) Representative whole-cell patch-clamp recordings (left panels) showing the effects of empagliflozin on sodium currents from single class III mutated HEK293T cells (centre panels). Grouped data showing the effects of empagliflozin on channel activation and inactivation (right panels). *Grouped data are from 5-10 cell recordings per group. Data are presented as means ± SEM; *p<0.05; **p<0.01; ***p<0.001. I/Imax indicates test current/maximum current; G/Gmax indicates test conductance/maximum conductance*.

**Figure 5.**
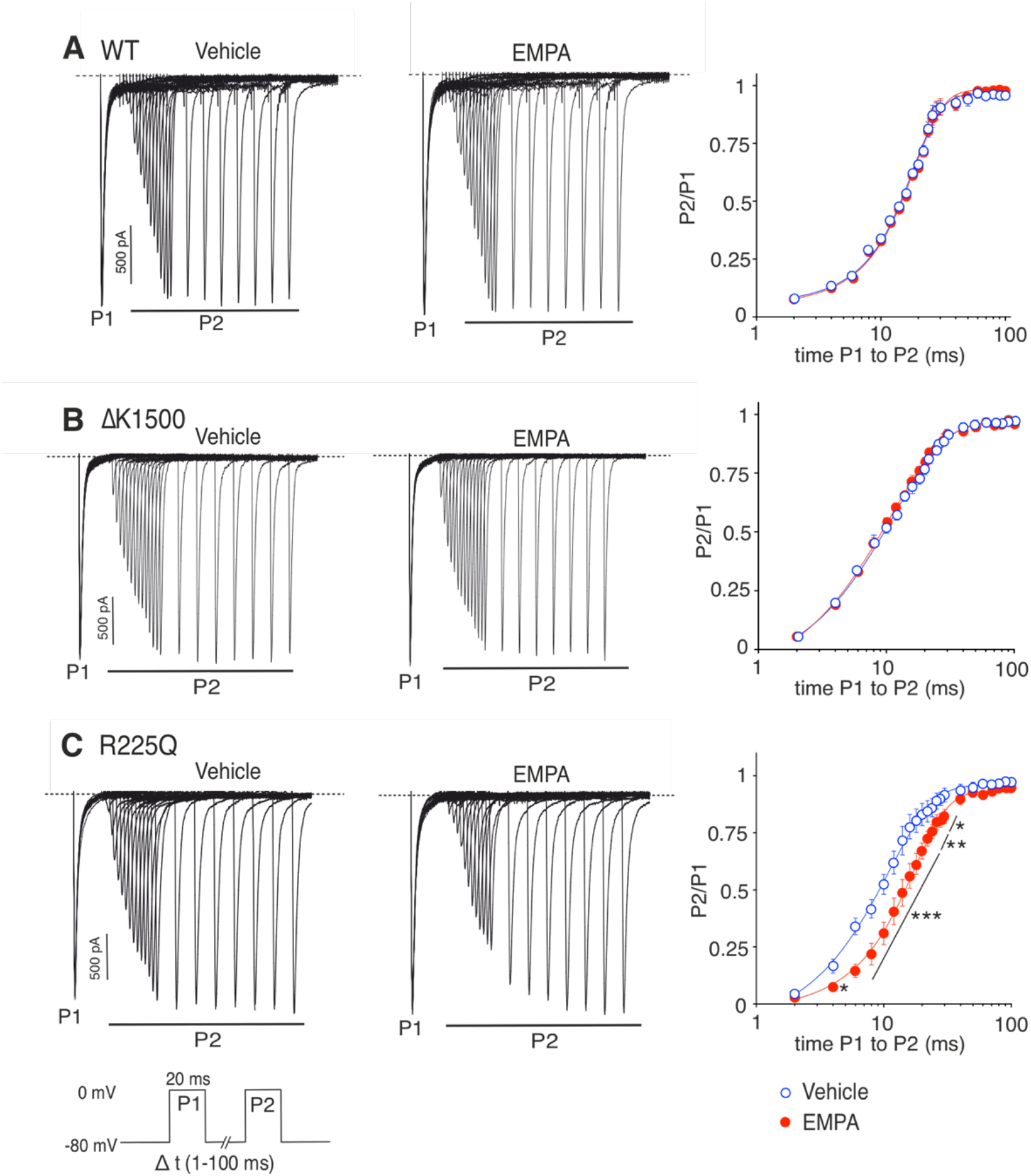
Empagliflozin does not slow recovery from inactivation in a long QT syndrome type 3 mutation present in the inactivation gate. Representative whole-cell patch-clamp recordings showing the effects of empagliflozin on time-dependent recovery from inactivation in single wildtype (A), class I representative ΔK1500 (B), and class II representative R225Q (C) mutated HEK293T cells, and time course of recovery from inactivation. *Fractional recovery was calculated by dividing the current magnitude of P2 by the corresponding P1. Curve was fitted with sigmoidal Boltzmann equation on a logarithmic scale. Data are presented as means ± SEM; *p<0.05; **p<0.01; ***p<0.001*.

### Implications for Ventricular Myocyte Excitation-Contraction Coupling

Our results on recombinantly expressed Nav1.5 containing LQT3 mutations provide initial support for putative therapeutic efficacy of empagliflozin in certain LQT3 mutations. However, it is important to integrate these findings with human ventricular myocyte excitation-contraction coupling. Access to primary myocytes from patients with LQT is not feasible, therefore we employed a mathematical model of the human ventricular AP and calcium handling. This computational approach offers an accurate means for directly representing experimentally measured properties of Nav1.5 gating kinetics in the context of myocyte electrophysiology. Experimental data on ΔKPQ, R225Q, and ΔK1500 from this study were first used to refit the Clancy *et al.* Markov model of cardiac I_Na_ that was originally developed to model LQT3 mutant gating [29; 56]. A key assumption of this model for ventricular myocytes is that late I_Na_ results from channels operating in a non-canonical burst mode (LC3>LO), which is a non-inactivating mode of gating (Figure 6I). A small fraction of channels transition into burst mode depending on a voltage-dependent transition rate (a8) and return to normal gating via b8. The refitting procedure (see supplemental materials) yielded 5 representative models (WT, Class I, Class II, Class I + empa, and Class II + empa) capturing the differences in steady-state properties, late I_Na_, and recovery from inactivation described above. Consistent with our experimental data, late I_Na_ was only selectively inhibited in ΔKPQ (class I) mutants (Figure 6A), and only Class II mutant R225Q demonstrated slowing of recovery (figure 6H). Simulation of recovery from inactivation for ΔK1500 was generated as a representative of a Class I mutation as we held extensive experimental recovery data of this mutant (Figure 6G). ΔKPQ activation and inactivation were unaltered by empagliflozin (Figure 6B), whereas R225Q had a negative shift of inactivation and positive shift of activation (Figure 6E).

**Figure 6.**
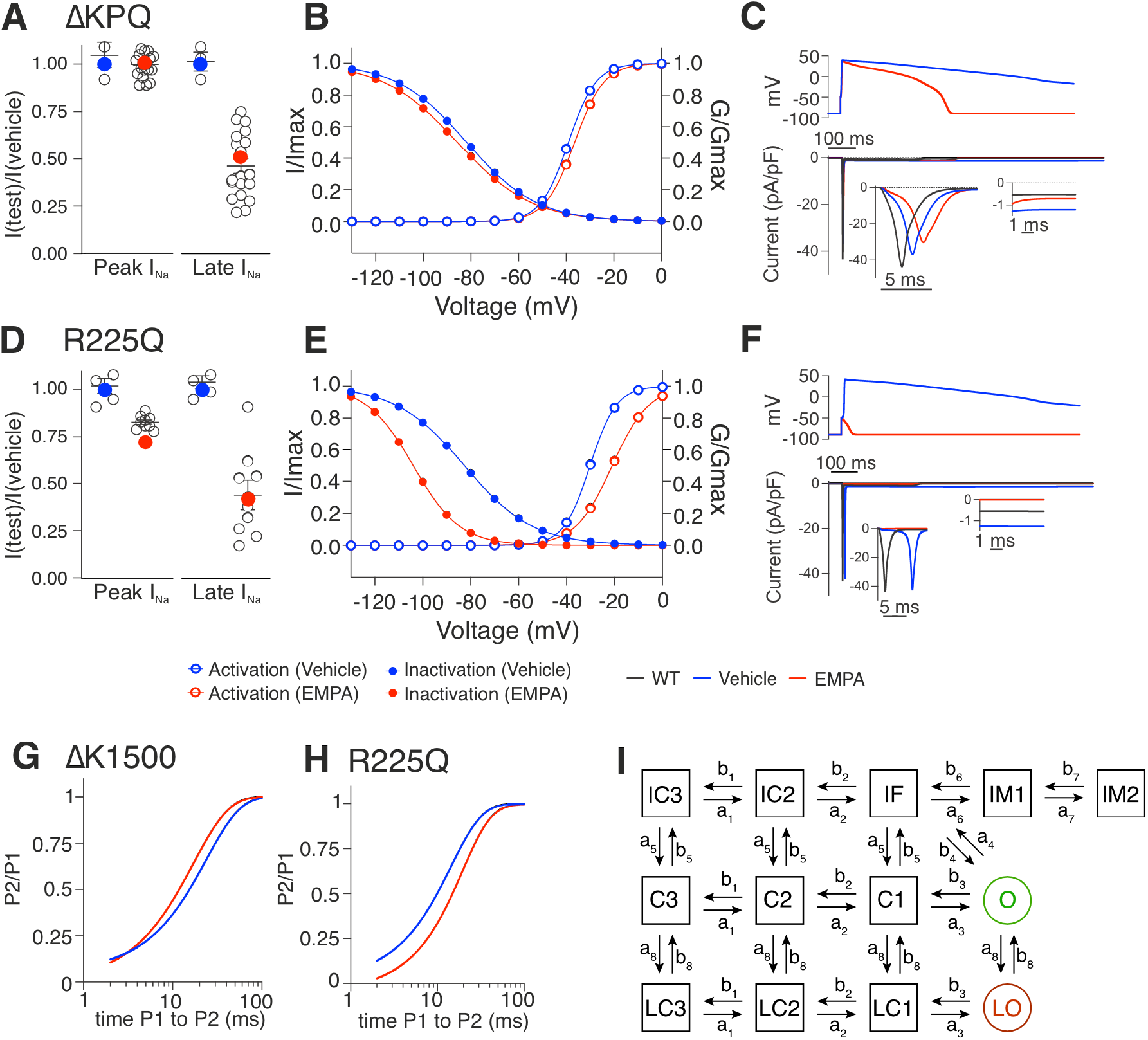
Computational model to determine the effect of empagliflozin on electrophysiology of ΔKPQ and R225Q mutant Nav1.5 channels in ventricular myocytes. Simulated data of peak and late INa in the class I, ΔKPQ (A) and class II, R225Q (D) mutant Nav1.5 models with/without (red/blue circles) 10 µM empagliflozin (EMPA). Data is illustrated as a ratio of test/vehicle current and open circles represent experimental data. Simulated steady-state properties of the ΔKPQ (B) and R225Q (E) models. Activation curves are constructed by I/Imax (test current/maximum current), and inactivation curves are constructed by G/Gmax (test conductance/maximum conductance). (C and F) Simulated upstroke and early repolarization (top), and corresponding INa (bottom, with zoomed insets of peak and late currents) for ΔKPQ (class I, C), and R225Q (class II, F). Current densities are normalized to capacitance (pA per pF). Simulated recovery from inactivation in ΔK1500 (G) and R225Q (H) channels is expressed as P2/P1 (dividing the current magnitude of P2 by the corresponding P1). (I) The Markov model structure of Nav1.5 gating published by Clancy *et al.*, with inactivated states (top), closed states (middle), and burst mode states (bottom) [29]. The open (O) and late open (LO) states are highlighted in green and red, respectively.

Each of these 5 I_Na_ models was then embedded into the Tomek-Rodriguez updates to the well-established O’Hara-Rudy human epicardial ventricular myocyte model (ToR-ORd) [30; 57]. In line with our experiments, as well as clinical manifestations, ΔKPQ and R225Q displayed prolonged AP duration and EADs (Figure 7B and 7C). EADs are the result of interrupted repolarization, resulting in premature influx of calcium ions via reactivation of L-type calcium current and sodium-calcium exchange [58]. Simulating the effects of empagliflozin unmasked normalization of the AP duration and elimination of EADs only in ΔKPQ. Simulating the effect of empagliflozin in R225Q resulted in a complete cessation of AP firing (Figure 7C), even after the I_Na_ conductance parameter (g_Na_) for this model was doubled compared to WT. As shown from our experimental data, empagliflozin reduces peak I_Na_ in R225Q channels and likely favors the inactivated state significantly enough to prevent regenerative I_Na_ activation in this model.

**Figure 7.**
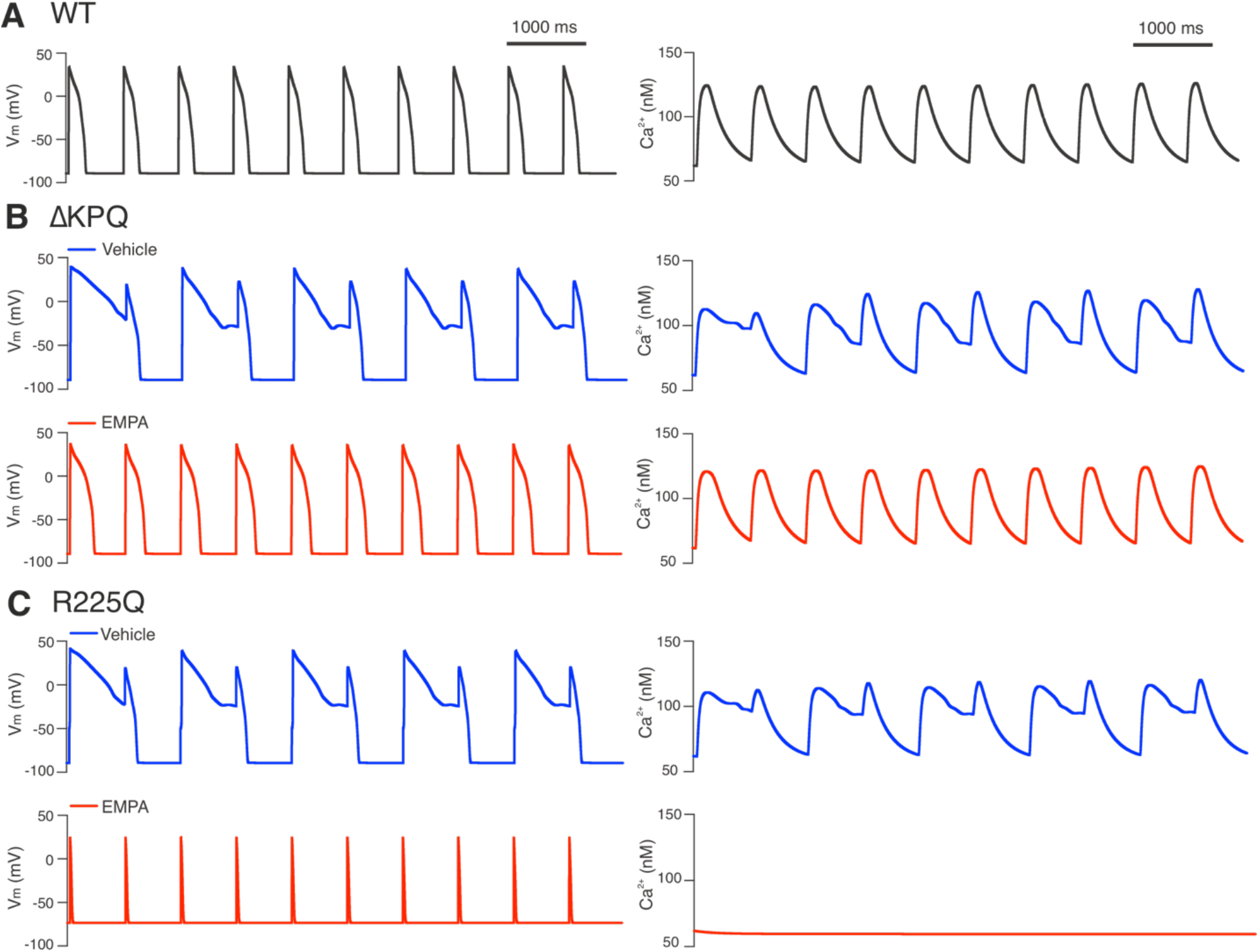
Early afterdepolarizations (EADs) during pacing in the class I (ΔKPQ) and class II (R225Q) models. Simulated action potentials when WT (A), ΔKPQ (B), and R225Q (C) Nav1.5 models are embedded in the ToR-ORd ventricular myocyte model. Action potential waveforms are shown at left and cytosolic calcium at right. Pacing was performed at 1 Hz. 10 µM empagliflozin is simulated for the two mutant models and restores the ΔKPQ mutant to stable repolarization but fails to elicit regenerative INa in the R225Q mutant despite ample stimulus. *VM = membrane potential*.

## Discussion

Results from this study revealed selective late I_Na_ inhibition by empagliflozin in HEK cells transfected with Nav1.5 channels containing known LQT3 mutations in the inactivation gate. Furthermore, the inhibitory effect of empagliflozin was selective to late I_Na_, sparing peak I_Na_ and recovery from inactivation. This late I_Na_ selectivity and lack of effect on gating kinetics were not observed in LQT3 mutants residing in the voltage-sensing regions (where peak current and kinetics were affected) nor or putative empagliflozin binding regions (where there was no late I_Na_ inhibition).

Excessive levels of late I_Na_ are associated with LQT3 syndrome, as well as in heart failure and other arrhythmias [59–61]. Inhibitors of late I_Na_ are therefore promising agents for treatment of heart disease and arrhythmias. Our previous research showed that empagliflozin selectively inhibits late I_Na_ in a transverse aortic constriction mouse model of heart failure [27]. Furthermore, empagliflozin reduced late I_Na_ in LQT3 mutants ΔKPQ and R1623Q, albeit not selectively in the latter mutation. Patients with LQT3 mutations have gain-of-function mutations of Nav1.5-encoding *SCN5A*, resulting in increased AP duration, detectable as prolonged QT interval on ECG. LQT3 patients have an increased risk of early and delayed afterdepolarizations, and life-threatening Torsade de Pointes-induced ventricular fibrillation. Inhibition of late I_Na_ reduces the risk of potentially fatal arrhythmias by reversing the prolonged duration of AP.

As our previous research hinted at differential mutant-specific late I_Na_ inhibition with empagliflozin in two mutants [27], we comprehensively tested the *in vitro* effects of empagliflozin on late I_Na_ in 13 previously identified LQT3 mutations residing in different locations of the channel. By investigating the extent of selective late I_Na_ inhibition by empagliflozin in many different LQT3 mutant sodium channels we aimed to shed new light on potentially novel precision treatment options for genetically screened patients with LQT3 syndrome. Based on location within the channel, mutations were classified into three Classes: I) the inactivation gate region, II) the S4 voltage-sensing regions, and III) the putative empagliflozin binding region. Our results demonstrate a selective inhibition of late I_Na_ only in Class I mutations. In Class II mutations, a concomitant inhibition of peak I_Na_ was observed, whereas Class III mutations did not show any inhibitory effect on neither late nor peak I_Na_. We speculate that empagliflozin may stabilize the inactivated state of Class II mutated Nav1.5 channels. As a result, slow recovery from inactivation ultimately represses peak I_Na_ of subsequent depolarizations. Therefore, we performed experiments to investigate time-dependent recovery from inactivation. As expected, empagliflozin significantly slowed recovery from inactivation in R225Q, a Class II mutant, whilst no change was observed in WT or the Class I mutant ΔK1500.

Interestingly, the absence of late I_Na_ inhibition in mutations of the putative empagliflozin binding regions (Class III), fDI-DIV and fDIII-DIV, suggests their involvement with empagliflozin binding. These regions are known binding sites for local anaesthetics like lidocaine, which is a selective inhibitor of open sodium channels, such as late I_Na_ or rapidly opening peak I_Na_. As a result, peak I_Na_ is inhibited in a frequency-dependent manner (i.e., more effective peak I_Na_ inhibition at higher heart rates), which was not observed with empagliflozin [27]. Although the exact binding site of empagliflozin remains to be mapped extensively, our results corroborate previously predicted empagliflozin binding sites within Nav1.5.

Previous research suggests an alternative explanation for the inhibitory effect on late I_Na_ via inhibiting CaMKII activity [26; 62]. Thus, inhibition of CaMKII by empagliflozin could indirectly inhibit late I_Na_ by reducing phosphorylation-dependent induction of late I_Na_. However, our results show that Class III LQT3 mutations located in the putative empagliflozin binding region result in a loss of late I_Na_ inhibition by empagliflozin. These results strongly support the concept of a direct inhibitory effect of empagliflozin on the channel itself rather than by any indirect mechanism. Therefore, it would be of interest to investigate the exact moieties involved in selective late I_Na_-inhibitory potency, to expedite development of specific late I_Na_ inhibitors without canonical SGLT2 inhibition in the renal proximal tubules.

Ideally, access to human cardiac tissue samples from patients with specific LQT3 mutations would allow a further assessment on the translatability of our results, although this is simply not possible. Therefore, to provide an alternative approach to extend our findings to human cardiac electrophysiology, simulations of the electrophysiological properties were generated in a mathematical model of human ventricular myocytes. Our experimental data was embedded sequentially in the Clancy I_Na_ and ToR-ORd models, and the resulting AP simulations were remarkably consistent with known clinical outcomes associated with these mutants. Results from these studies extend of mechanistic findings by demonstrating a favourable electrophysiological response to empagliflozin in simulations where a Class I LQT3 mutation is modelled.

Low micromolar concentrations of empagliflozin used in this study correspond to the steady-state range in human patients [63]. Therefore, the results from our study may be directly translatable to a clinical setting in which patients with certain LQT3 mutations may be treated with SGLT2is within the well-established therapeutic range for this class of drug. Compared to the current restricted treatment options for patients with LQT3 mutations, empagliflozin may represent a novel orally bioavailable and non-invasive alternative with few side effects. Future studies should include 1) cellular studies on gene-edited stem cells differentiated into cardiac cells and 2) clinical studies on LQT3 patients harbouring *SCN5A* Class I mutations to determine the effects of empagliflozin on ECG QT intervals.

In summary, our findings also provide a promising novel precision medicine approach for patients with certain LQT3 mutations in which empagliflozin selectively inhibits late I_Na_.

## Acknowledgments

This research was supported by a project grant to P.E.L. from the Canadian Institutes of Health Research. L.L. performed experiments, analysed data, generated figures and co-wrote the manuscript. M.F., W.L., A.B. and B.G. performed experiments, generated the mutations, analysed the data, assembled figures and contributed to manuscript writing. A.C and K.B. performed all of the in silico structural modelling and contributed to manuscript writing. A.G.B. performed all of the mathematical modelling studies and contributed to manuscript writing. P.E.L. conceived the original concept of the project, contributed to all aspects of the experimental design, data analysis, interpretation of the results, generation of the figures and is the senior author of the manuscript.

## S1 Supplementary Methods

### Mathematical Models of Nav1.5 Gating

We developed the five models of I_Na_ (WT, ΔKPQ, R225Q, ΔKPQ+empa, and R225Q+empa) from the 13-state Markov model of cardiac I_Na_ originally published by Clancy et al. [1]. This model is shown in main Figure 6I. It assumes that late I_Na_ is an alternative gating mode of Nav1.5, and thus directly couples the dynamics of normal fast I_Na_ gating to late I_Na_ dynamics. The late or “burst” mode states are LC3 – LO, where only LO is open and available to conduct current. Transitions from C3 – O represent canonical voltage dependent activation of fast I_Na_, and in reverse, deactivation. States IC3 – IM2 represent inactivated states, where transition from O to IF is canonical fast inactivation, and IM1 and IM2 represent more deeply inactivated states with longer recover times. Transitions in the direction from IM2 – IC3 determine the various temporal components of recovery from inactivation.

To recapitulate the Nav1.5 dynamics observed in this study, and previously in Philippaert et al. [2], we reformulated and refitted the Clancy model. To permit differing I_Na_ slope-factors for steady-state activation and inactivation in all, we first reformulated the a5 transition rate (from Fig 6I) to take the general form:

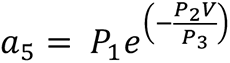

We then refit WT model parameters to: (1) steady-state activation and inactivation (Fig 1G in [2]); (2) percent late I_Na_ (Fig 1A in [2]); (3) recovery from inactivation (data herein, Figure 5). This WT model served as the starting point for fitting both mutant models (ΔKPQ and R225Q), again for steady-state properties, late I_Na_ and recovery form inactivation. Finally, to generate the ΔKPQ+empa and R225Q+empa models we fitted each of those mutant models for empagliflozin effects where they were observed. The resulting parameter sets are provided in Table S1.

In all cases, parameter fitting was performed via the Nelder-Mead simplex optimization algorithm fminsearch in Matlab (Release 2022b, Mathworks, Natick MA). Parameters for fitting steady state properties were constrained to include only those impacting rates a_1_ to a_5_. Parameters for fitting late I_Na_ were constrained to include only a_8_ and b_8_. For recovery from inactivation, only rates a_6_ to a_7_ and b_6_ to b_7_ were allowed to vary.

### Human Ventricular Myocyte Action Potential Model (ToR-ORd)

Upon substituting the WT I_Na_ model derived above into the ToR-ORd action potential model (epicardial model version), repolarization was unstable due to large I_Na_ currents. Thus, we reduced whole-cell I_Na_ conductance (g_Na_) to stability at 3.0 mS/μF. All other parameters were unchanged. For the R225Q model, due to marked left-shifted steady-state inactivation, this resulted in silencing of the myocyte model. Thus, to observe the effects of empagliflozin on the paced phenotype, we doubled WT g_Na_ to 6.0 for both the R225Q and R225Q+empa simulations. For consistency across simulations, we paced ToR-Ord models with 0.5 ms square current injections of -80 pA/pF, and at 1 Hz.

Final code for each of the Nav1.5 models and final ToR-Ord simulations is accessible at https://github.com/andygedwards/LQT3-SGLT2i

**Table S1.** Parameters for NaV1.5 models.

| Transition rate | Parameter | Models |  |  |  |  |
| --- | --- | --- | --- | --- | --- | --- |
| | | WT | ΔKPQ | R225Q | ΔKPQ+empa | R225Q+empa |
| $a_1$ | $a_1P1$ | 3.802 | 3.802 | 3.802 | 3.802 | 3.802 |
| | $a_1P2$ | 0.1027 | 0.1027 | 0.1027 | 0.1027 | 0.1027 |
| | $a_1P3$ | 12.83 | 11.62 | 9.62 | 10.88 | 28.33 |
| | $a_1P4$ | 0.1433 | 0.145 | 0.1428 | 0.1238 | 0.3058 |
| | $a_1P5$ | 179.3 | 182.1 | 186.2 | 248.9 | 15.63 |
| $a_2$ | $a_2P1$ | 3.802 | 3.802 | 3.802 | 3.802 | 3.802 |
| | $a_2P2$ | 0.1027 | 0.1027 | 0.1027 | 0.1027 | 0.1027 |
| | $a_2P3$ | 18.75 | 17.23 | 10.37 | 17.46 | 19.47 |
| | $a_2P4$ | 0.2727 | 0.2702 | 0.309 | 0.2633 | 0 |
| | $a_2P5$ | 90.26 | 88.12 | 79.89 | 75.19 | N/A |
| $a_3$ | $a_3P1$ | 3.802 | 3.802 | 3.802 | 3.802 | 3.802 |
| | $a_3P2$ | 0.1027 | 0.1027 | 0.1027 | 0.1027 | 0.1027 |
| | $a_3P3$ | 35.51 | 35 | 36.72 | 30.29 | 10.81 |
| | $a_3P4$ | 0.2106 | 0.2499 | 0.241 | 0.2172 | 0.4673 |
| | $a_3P5$ | 35.09 | 33.75 | 29.73 | 27.87 | 13.17 |
| $b_1$ | $b_1P1$ | 0.1917 | 0.1917 | 0.1917 | 0.1917 | 0.1917 |
| | $b_1P2$ | 20.3 | 20.3 | 20.3 | 20.3 | 20.3 |
| $b_2$ | $b_2P1$ | 0.2 | 0.2 | 0.2 | 0.2 | 0.2 |
| | $b_2P2$ | -5 | -5 | -5 | -5 | -5 |
| | $b_2P3$ | 20.3 | 20.3 | 20.3 | 20.3 | 20.3 |
| $b_3$ | $b_3P1$ | 0.22 | 0.22 | 0.22 | 0.22 | 0.22 |
| | $b_3P2$ | -10 | -10 | -10 | -10 | -10 |
| | $b_3P3$ | 20.3 | 20.3 | 20.3 | 20.3 | 20.3 |
| $a_4$ | $a_4P1$ | 9.178 | 9.178 | 9.178 | 9.178 | 9.178 |
| | $a_4P2$ | 29.68 | 29.68 | 29.68 | 29.68 | 29.68 |
| $b_4$ | N/A | $a_3 \cdot a_4 \cdot a_5 / (b_3 \cdot b_5)$ | $a_3 \cdot a_4 \cdot a_5 / (b_3 \cdot b_5)$ | $a_3 \cdot a_4 \cdot a_5 / (b_3 \cdot b_5)$ | $a_3 \cdot a_4 \cdot a_5 / (b_3 \cdot b_5)$ | $a_3 \cdot a_4 \cdot a_5 / (b_3 \cdot b_5)$ |
| $a_5$ | $a_5P1$ | 1.83E-04 | 2.23E-04 | 2.39E-04 | 3.23E-04 | 8.58E-04 |
| | $a_5P2$ | 0.3764 | 0.3744 | 0.3984 | 0.3799 | 0.499 |
| | $a_5P3$ | 5.442 | 5.94 | 5.829 | 6.251 | 4.863 |
| $b_5$ | $b_5P1$ | 0.0373 | 0.0397 | 0.0696 | 0.0599 | 0.377 |
| | $b_5P2$ | 2.00E-05 | 2.00E-05 | 2.00E-05 | 2.00E-05 | 4.19E-05 |
| $a_6$ | $a_6P1$ | 24.44 | 24.75 | 23.69 | 20.02 | 22.12 |
| $b_6$ | $b_6P1$ | 0.5033 | 0.5479 | 0.4756 | 0.5042 | 0.3531 |
| $a_7$ | $a_7P1$ | 2050 | 2377 | 2465 | 1830 | 225.7 |
| $b_7$ | $b_7P1$ | 0.213 | 0.2086 | 0.211 | 0.2289 | 0.3038 |
| $a_8$ | $a_8P1$ | 2.09E-05 | 2.47E-05 | 1.66E-05 | 2.29E-05 | 2.48E-05 |
| $b_8$ | $b_8P1$ | 2.90E-03 | 1.06E-03 | 1.27E-03 | 1.70E-03 | 2.98E-03 |

